# Characterization of Cas proteins for CRISPR-Cas editing in streptomycetes

**DOI:** 10.1101/526996

**Authors:** Wan Lin Yeo, Elena Heng, Lee Ling Tan, Yi Wee Lim, Yee Hwee Lim, Shawn Hoon, Huimin Zhao, Mingzi M. Zhang, Fong Tian Wong

**Affiliations:** Metabolic Engineering Research Laboratory, Institute of Chemical and Engineering Sciences, A*STAR, Singapore; Molecular Engineering Laboratory, Biomedical Institutes of Sciences, A*STAR, Singapore; Organic Chemistry, Institute of Chemical and Engineering Sciences, A*STAR, Singapore; Department of Chemical and Biomolecular engineering, Department of Chemistry, Department of Biochemistry, University of Illinois at Urbana-Champaign, United States; Institute of Molecular and Genomic Medicine, National Health Research Institutes, Taiwan, R.O.C.

**Keywords:** CRISPR-Cas, *Streptomyces*, genome editing

## Abstract

Application of the well-characterized *Streptococcus pyogenes* CRISPR-Cas9 system in actinomycetes has enabled high efficiency multiplex genome editing and CRISPRi-mediated transcriptional regulation in these prolific bioactive metabolite producers. Nonetheless, SpCas9 has its limitations and can be ineffective depending on the strains and target sites. Here, we built and tested alternative CRISPR-Cas constructs based on the standalone pCRISPomyces-2 editing plasmid. We showed that *Streptococcus thermophilus* CRISPR1 (sth1Cas9), *Staphylococcus aureus* Cas9 (saCas9), and *Francisella tularensis subsp. Novicida* U112 Cpf1 (fnCpf1) are functional in multiple streptomycetes enabling efficient homology directed repair (HDR)-mediated knock-in and deletion. In strains where spCas9 was nonfunctional, these alternative Cas systems enabled precise genomic modifications within biosynthetic gene clusters for the discovery, production and diversification of natural products. These additional Cas proteins provide us with the versatility to overcome the limitations of individual CRISPR-Cas systems for genome editing and transcriptional regulation of these industrially important bacteria.

## Introduction

The CRISPR-Cas technology has revolutionized biological research and the biotech industry. Since the first application of Class 2 CRISPR-Cas systems to edit genomes (Cong, *et al*, 2013), the number and diversity of CRISPR-Cas applications (Adli, 2018) have increased exponentially. The rapid adoption of this technology is due to its capability for precise editing and ease of use – a short RNA transcript is sufficient to guide the nuclease to its desired target, as well as the wide range of organisms that it can be applied in. These advantages have led us and other labs to explore CRISPR-Cas editing in streptomycetes (Li, *et al*, 2018, Zhang, *et al*, 2018, Tao, *et al*, 2018), which are prolific producers of bioactive secondary metabolites. Previously, we demonstrated that CRISPR-Cas increases the speed and throughput of genome editing compared to the traditional two step selection-counterselection protocol, allowing rapid genetic access to natural products encoded in the genomes of hitherto genetically intractable streptomycetes (Zhang, *et al*, 2018).

To date, *Streptomyces* strains have been mostly engineered using the well-characterized *Streptococcus pyogenes* Cas9 (spCas9, Tao, *et al*, 2018). Yet as we have observed in our labs and others (Moreb, *et al*, 2017, Cui, *et al*, 2018), a major bottleneck is that many bacteria strains still cannot be edited by *S. pyogenes* Cas9. In some examples, after a successful introduction of recombinant CRISPR-Cas systems, some bacteria can still evade modifications induced by CRISPR-Cas9 double stranded breaks with RecA-dependent repair (Moreb, *et al*, 2017). In other strains or even in certain target sites of previously editable strains, no exconjugants can be obtained in the presence of Cas9 and its guide RNA, possibly due to off-target nuclease-induced toxicity or tight binding of Cas9 to nontargets in the absence of guide RNAs (Cui, *et al*, 2018). To overcome similar bottlenecks in streptomycetes, we were motivated to expand the CRISPR-Cas toolbox. In this study, we compared different Class 2 Cas proteins and showed that these alternative Cas proteins allow us to overcome the bottlenecks faced with using spCas9 to facilitate natural product discovery in these industrially important bacteria.

## Materials and methods

### Reagents and Media

Unless otherwise indicated, all reagents are obtained from Sigma. 1 L of MGY medium contains 10 g malt extract broth, 4 g Bacto yeast extract (BD Biosciences), 4 g glucose (1st Base, Axil Scientific) and for MGY agar plates, an additional 20 g of Bacto agar (BD Biosciences). ISP Medium No. 4 agar (BD Biosciences) for *Streptomyces* sp. NRRL S-244. Conjugation experiments involving WM6026 and WM3780 *E. coli* strains were performed on R2 agar without sucrose: 0.25 g K_2_SO_4_, 10.12 g MgCl_2_•6H_2_O, 10 g glucose, 0.1 g Bacto casamino acids (BD Biosciences), 5.73 g TES, 20 g agar in 1 L water, autoclaved, after which 1 mL filter-sterilized 50 mg/mL KH_2_PO_4_ solution and filter-sterilized 2.94 g CaCl_2_•2H_2_O and 3 g L-proline in 5 mL 1 N NaOH were added to the medium.

### Strains and Growth conditions

Unless otherwise indicated, strains are propagated in MGY medium at 30 °C. Spore preparations and conjugation protocols were similar as described before (Aínsa, *et al*, 2000). For spore preparations, 1:1000 of a spore preparation or 1:100 dilution of a saturated seed culture is plated on MGY or ISP Medium No. 4 plates and incubated at 30 °C until thick spores are observed. Spores were removed from the plate using 5 mm glass beads (Sigma) and resuspended in sterile TX buffer (50 mM Tris pH 7.4, 0.001% (v/v) Triton X) by vigorous vortexing for 2 min. The suspension was then passed through a syringe containing a cotton ball. The eluant containing free spores were pelleted by spinning at maximum speed in an Eppendorf 5810R centrifuge for 10 min, resuspended in 1 mL sterile water and pelleted again. The spores were then resuspended in water and stored at −80 °C. A typical spore prep contains ~10^6^–10^7^ spores/mL as determined by serial dilution plating.

### Target site prediction

Target sites are predicted using modifications to PhytoCRISP-Ex software (Rastogi, et al, 2016). Potential sites are given as sites where seed regions (last 15 bases including PAM sequences) do not have 100% match with off-targets.

### Construction of genome editing plasmids

The different Cas proteins were codon-optimized based on the codon usage of *Streptomyces coelicolor* and synthesized by Genscript. All DNA manipulations were carried out in OmniMAX™ (Thermo Fisher). Primers used in this study are listed in Supplementary Table 1. Restriction enzymes were obtained from New England Biolabs. Protospacers were first inserted via BbsI-mediated Golden Gate Assembly before introduction of the respective homology flanks *via* Gibson assembly, as previously described (Zhang, *et al*, 2017).

### Interspecies conjugation

Promoter knock-in constructs were used to transform conjugating *E. coli* strains and colonies with apramycin resistance (50 mg/L) were picked into LB with apramycin. WM6026 requires diaminopimelic acid (0.3 mM) in LB for growth and it was added to LB for subsequent wash and resuspension steps. Overnight cultures were diluted 1:100 into fresh LB with antibiotics and grown to an OD_600_ of 0.4–0.6. 400 μL of the culture was pelleted, washed twice and resuspended in LB without antibiotics. The washed *E. coli* cells were then mixed with spores at 1:5 volume ratio and spotted on R2 without sucrose plates. After incubation for 16–20 h at 30 °C, the plates were flooded with nalidixic acid and apramycin and incubated until exconjugants appear. Exconjugants were streaked onto MGY or ISP Medium No. 4 plates containing apramycin at 30 °C followed by restreaking to MGY or ISP Medium No. 4 plates at 37 °C to cure the CRISPR-Cas9 plasmid containing a temperature-sensitive origin of replication. Apramycin-sensitive clones growing at 37 °C were then subjected to validation of promoter knock-in and genome editing as described below.

### Validation of promoter knock-in and genome editing

Genomic DNA from wild type and exconjugants from the indicated strains were isolated from liquid cultures using the Blood and Tissue DNeasy kit (Qiagen) after pretreating the cells with 20 mg/mL lysozyme for 0.5–1 h at 30 °C. PCR was performed using control primers beyond the homology regions (Supplementary Table 1) with KODXtreme Taq polymerase (Millipore). Where indicated, PCR products were subjected to digest with specific restriction enzymes to differentiate between PCR products of wild type genomic sequences and successful genome editing by knock-ins. Positive samples were purified using ExoSAP-IT™ (Affymetrix USB) and validated by Sanger sequencing (Figure S1, S2).

### Fermentation, extraction and NMR of phosphonate compound

Liquid seed cultures (2 mL MGY) of wild type and engineered S. sp. NRRL S-244 strains were inoculated from a plate or spore stock in 14 mL culture tubes. Seed cultures were incubated at 30 °C with 250 rpm shaking until achieving turbidity or high particle density (2 days). Seed cultures were diluted 1:100 into 50 mL of MGY broth in 250 mL baffled flasks containing ~30–40 5 mm glass beads and incubated at 30 °C with 250 rpm shaking for 7 days. The cultures were harvested by pelleting at maximum speed in an Eppendorf 5810R centrifuge for 10 min. The supernatants were split into two 50 mL falcon tubes. Culture supernatants were lyophilized and then extracted with equal volume of methanol, before evaporated to dryness.

The extracts were dissolved in D_2_O and transferred to a 5 mm NMR tube for NMR analysis. ^31^P-NMR has been acquired using a Bruker DRX-400 spectrometer. Proton decoupled ^31^P-NMR spectra are referenced to an external H_3_PO_4_ (aq) standard (δ 0.0 ppm). All samples have been acquired for 2000 scans. Comparison of production titer of various strains was done by spiking in known amounts of authentic FR-900098 (Sigma) as reference standard.

## Results

### Alternative Cas proteins for genome editing of streptomycetes

In this study, we tested three Cas proteins in addition to our original spCas9 construct for genome editing in streptomycetes (**Figure 1**). The Cas proteins include Type II Cas from *S. pyogenes* (spCas9), *Streptococcus thermophilus* CRISPR1 (sth1Cas9, Horvath, *et al*, 2008), *Staphylococcus aureus* (saCas9, Ran, *et al*, 2015) and *Francisella tularensis subsp. Novicida U112’s* Type V-A Cas (fnCpf1, Zetsche, *et al*, 2015). Sth1Cas9 and saCas9 are Type II Cas proteins with similar editing efficiencies but are about 25% smaller in size compared to spCas9 (Horvath, *et al*, 2008; Ran, *et al*, 2015). fnCpf1 is a Type V-A single-effector nuclease that is closer in size to spCas9 (Zetsche, *et al*, 2015). Both Class 2 CRISPR-Cas systems utilize RNA guides to target double stranded DNA with several key differences (Zetsche, *et al*, 2017, **Figure 1**). Type II results in a blunt cut whilst Type V-A yields a staggered cut that is preferential towards homology directed repair (HDR) instead of the error-prone non-homologous end-joining (NHEJ) (**Figure 1A**). Unlike Type II Cas proteins, Type V-A proteins do not require a tracrRNA and a separate RNA processing protein since pre-crRNA can be wholly processed by the nuclease effector. This difference in crRNA processing for Type V-A class proteins allow multiple targets to be transcribed on a single transcript for more rapid and facile multiplexing (Cobb, *et al*, 2014). The protospacer adjacent motifs (PAMs) for Type II proteins are also more GC-rich compared to the AT-rich PAMs for Type V-A (**Figure 1B**). Due to GC-rich streptomycete genomes, spCas9 provides the highest number of targets compared to the other Cas proteins (**Table 1**), but have proven ineffective in certain contexts such as those described later in this study. In general, we expect the use of different Cas proteins with distinct PAM requirements to increase our target density, which would be advantageous when targeting short intergenic regions of the genome.

**Figure 1.**
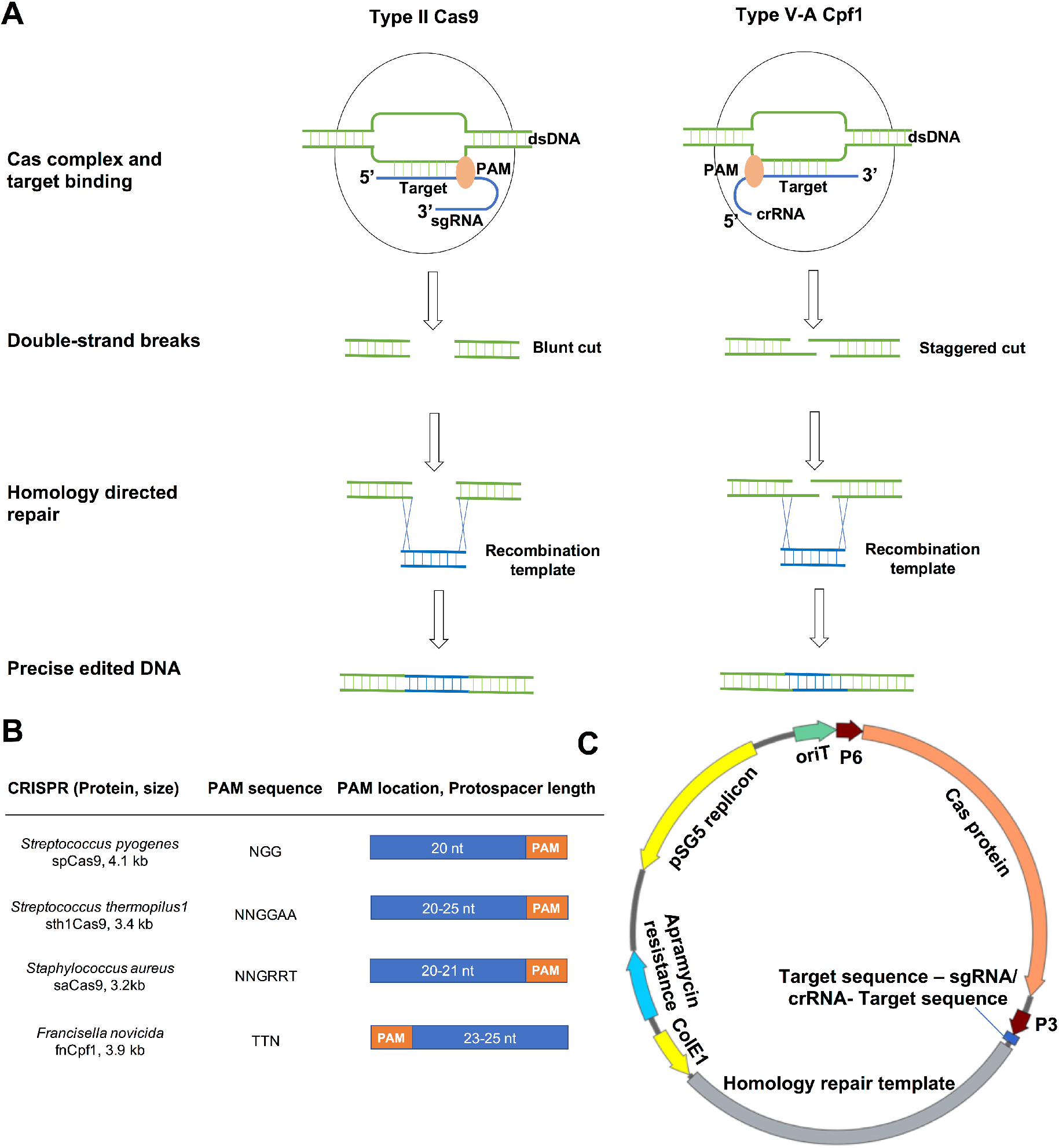
CRISPR-Cas systems used in this study. (A) Comparison of Type II and Type V-A Class 2 Cas systems. (B) Comparison of different Cas proteins and their target sequence requirements. (C) General map of all-in-one editing CRISPR-Cas constructs for one-step genome editing of streptomycetes. PAM, protospacer adjacent motif. P3 and P6 are promoters.

**Table 1.** *In silico* analyses of potential target sites for the indicated Cas proteins and their respective preferred PAMs. Length of target sequence for each case is indicated in brackets. The most preferred PAM sequences for the respective Cas9 proteins were used for this analysis.

| **Cas/PAM** | **Potential target sites** |  |  |
| --- | --- | --- | --- |
|  | ***Streptomyces albus***<br>**J1074 (3.8 kb)** | ***Streptomyces* sp.**<br>**NRRL S-244 (2.0 kb)** | ***Streptomyces roseosporus***<br>**NRRL 15998 (5.7 kb)** |
| **spCas/NGG** | 228 | 109 | 327 |
| **saCas9/NNGRRT** | 54 | 32 | 85 |
| **sth1Cas9/NNAGAA** | 9 | 3 | 19 |
| **fnCpf1/TTN** | 63 | 29 | 117 |

To test the utility of these different Cas proteins in different *Streptomyces* strains, we chose to focus on the activation and engineering of biosynthetic gene clusters (BGCs), which involve the insertion of small promoter elements into short intergenic regions and genetic manipulation of single genes. The Cas constructs were based on the all-in-one pCRISPomyces-2 editing plasmid (Addgene #61737, Cobb, *et al*, 2014) with the respective Cas protein and sgRNA/crRNA under the control of the two distinct promoters (**Figure 1C**). The final editing constructs was constructed by Golden Gate assembly to insert the targeting guide sequences, followed by the addition of homology repair templates at the XbaI site using isothermal assembly as previously described (Zhang, *et al*, 2017). We sought to target the different Cas proteins to the same genomic region as much as possible, but this is not always possible due to differences in PAM sequence requirements. Choices of target sites and homology repair templates were further limited by the short sequence targeted and our decision to minimize disruption to native BGC sequences. While differences in repair templates are mostly minimal and not expected to have significant impact on editing efficiencies, we acknowledge that selection of target sites is important for editing efficiency and should be considered when comparing the different Cas proteins.

### Cas proteins are functional in *Streptomyces albus*

We first validated the functionality of the different Cas proteins for genome editing in *Streptomyces albus*, a model streptomycete with high conjugation efficiencies. Target sites for all four Cas proteins can be found in the 280 bp target intergenic region of the indigoidine cluster (**Figure 2**). Consistent with our previous study (Zhang, *et al*, 2017), knock-in efficiency of the *kasO\** promoter was increased when spCas9 was targeted by the sgRNA to the region upstream of gene encoding for indigoidine synthase compared to the control where no guide was provided (100% vs 37%). The other three Cas proteins also yielded high editing efficiencies of 87-100% as validated by sanger sequencing (**Table 2**). These results demonstrate that the different Cas proteins are functional in *S. albus* and may be used in place of spCas9 for genome editing of *Streptomyces* strains.

**Figure 2.**
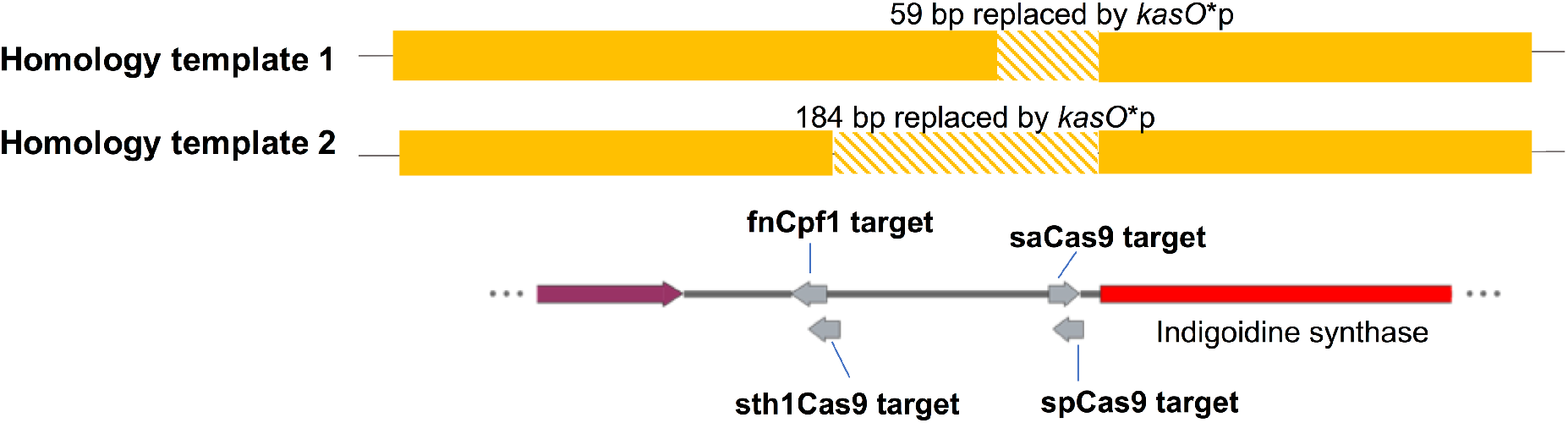
Target sequences for the different Cas proteins in *S. albus* for insertion of *kasO*\*p in front of indigoidine synthase. Intergenic region between the two genes are 280 bp. Homology arms of the editing templates (solid yellow) are aligned to the scheme of the genome and are designed to minimize disruption to native genome. For sth1Cas9, since NNAGAA is not available with the target sequence, the protospacer for sth1Cas9 was designed using an alternative NNGGAA PAM (Jiang, *et al*, 2017).

**Table 2.** Editing efficiencies and exconjugant output from Cas experiments.

| **Cas constructs** | **Editing efficiency<sup>†</sup>** | **# exconjugants<sup>‡</sup>** |
| --- | --- | --- |
| ***Streptomyces albus* J1074** |  |  |
| No protospacer control (spCas9, template 1) | 37 % (3/8) | >400 |
| spCas9 (template 1) | 100 % (8/8) | >400 |
| saCas9 (template 1) | 87 % (7/8) | >400 |
| sth1Cas9 (template 2) | 100 % (8/8) | >400 |
| fnCpf1 (template 2) | 87 % (7/8) | >400 |
| ***Streptomyces* sp. NRRL S-244** |  |  |
| No protospacer control (spCas9) | 0 % (0/8) | 29 |
| spCas9 | No exconjugants | 0 |
| saCas9 | 100 % (3/3) | 3 |
| fnCpf1 | 100 % (8/8) | 38 |
| ***Streptomyces roseosporus* NRRL 15998** |  |  |
| No protospacer control (spCas9) | 12.5 % (1/8) | 89 |
| spCas9 (targets 1-3) <sup>§</sup> | No exconjugants | 0 |
| saCas9 (target 1) | 100 % (3/3) | 3 |
| saCas9 (target 2) | 100 % (2/2) | 2 |

<sup>†</sup> Editing efficiency given by the number of correct clones validated by sanger sequencing over the total number of clones picked.
<sup>‡</sup> Number of exconjugants observed per 100 μL of spore preparation used in each conjugation. A typical spore prep contains $\sim 10^6$ - $10^7$ spores/ mL as determined by serial dilution plating.
<sup>§</sup> In multiple independent experiments with spCas9 targeting three different sites, no exconjugants were observed.

### Alternative Cas proteins enable genome editing in previously inaccessible contexts

After validating that the codon-optimized Cas proteins are functional and can be used in place of spCas9 to increase promoter knock-in efficiency in *S. albus*, we next examined how these Cas proteins will perform in applications where spCas9 has failed. Previous attempts using spCas9 to introduce a strong constitutive *kasO\** promoter in front of a putative *Streptomyces* antibiotic regulatory protein (SARP) to increase production of a low-yielding phosphonate compound in *Streptomyces* sp. NRRL S-244 had failed with no observed exconjugants (**Figure 3A, Table 2**). Highlighting the need for multiple Cas proteins, we were unable to find target sequences for sth1Cas9 (PAM NNAGAA) within range of the 64 bp target intergenic region. Nonetheless with saCas9, we managed to obtain a small number of exconjugants were observed with 100% editing efficiency. With fnCpf1, more exconjugants were observed with 100% editing efficiency (**Table 2**). In the end, a 10-fold upregulation in phosphonate production was also observed and validated in shake flask fermentation of two edited strains (**Figure 3C**).

**Figure 3.**
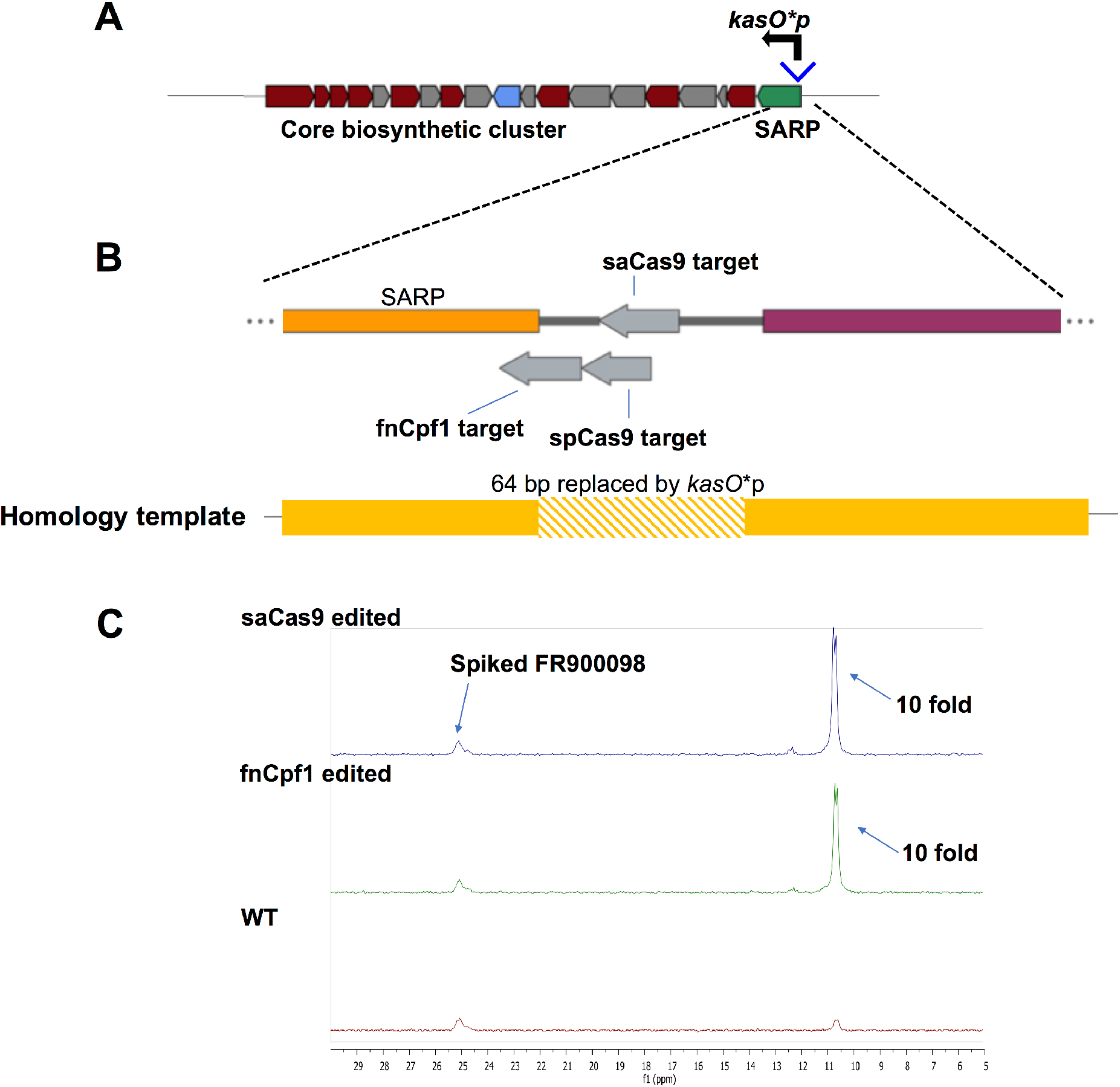
Upregulation of phosphonate production in *Streptomyces* sp. NRRL S-244. (A) Introduction of *kasO*\*p in front of SARP within the putative phosphonate BGC as annotated by AntiSMASH (Weber, *et al*, 2015). Red encodes biosynthetic enzymes, blue for transport protein and green for regulatory proteins. (B) Positions of target sequences for indicated Cas proteins. Homology arms of the editing templates (solid yellow) are aligned to the scheme of the genome. (C) ^31^P-NMR for two edited clones in comparison to wild type strain. Spiked FR-900098 is used as an internal standard for fold change calculations.

In another example, spCas9 has worked robustly in *Streptomyces roseosporus* for both insertions and deletions for multiple BGCs (Zhang, *et al*, 2017, Lim, *et al*, 2018), but has failed in a specific context. Attempts to delete a specific glycosyltransferase *aurS5* gene in a polyene macrolactam BGC (Lim, *et al*, 2018) yielded no exconjugants in the presence of spCas9 and a variety of guide RNAs targeting different locations (**Figure 4**). Interestingly, in the absence of a functional guide, edits could be observed, possibly a result of innate homologous recombination in the presence of the editing template (**Table 2**). In this case, alternative Cas proteins circumvented this problem with 100% editing efficiency observed for two saCas9 targets (**Figure 4, Table 2**).

**Figure 4.**
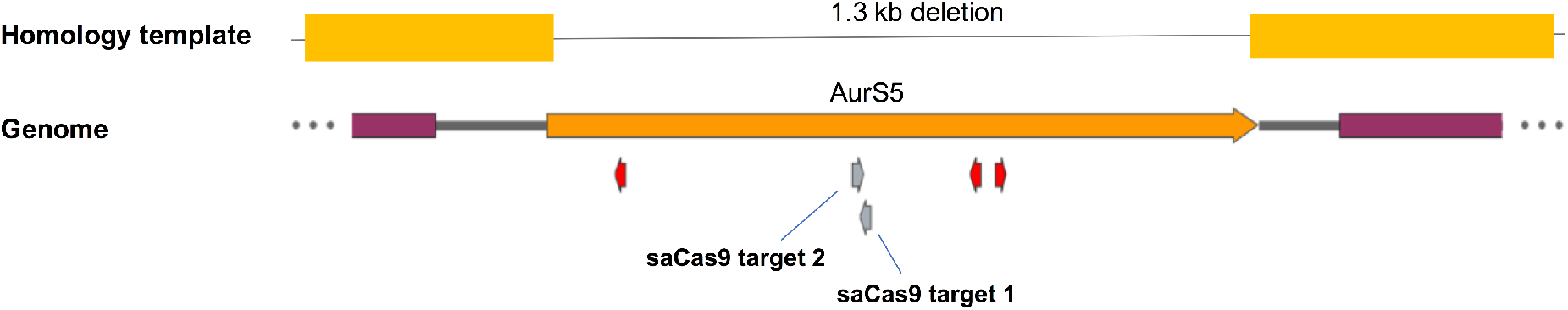
Scheme of targets and homology repair template used for *S. roseosporus* deletion. Red and grey arrows indicate the locations of spCas9 and saCas9 targets tested respectively.

## Discussion

Here, we present alternative CRISPR-Cas systems for genome editing in streptomycetes and showcase their potential to circumvent some of the limitations associated with the widely used and well characterized *S. pyogenes* CRISPR-Cas system (Zhang, *et al*, 2018, Tao, *et al*, 2018). While Cpf1 systems have recently examined for streptomycetes (Li, *et al*, 2018), this would be the first application of sth1Cas9 and saCas9 systems in streptomycetes. While the different Cas proteins all work in the model streptomycete *S. albus*, the alternative Cas toolbox proved to be useful in achieving precise genomic modifications within biosynthetic gene clusters for the discovery, production and diversification of natural products in contexts where spCas9 has failed. We also expect this expanded Cas toolbox to be relevant to the engineering of rare actinomycetes (Liu, *et al*, 2018). With these observations, this study also provides the motivation for systematic evaluation of different CRISPR-Cas systems in the genetic editing of *Streptomyces* strains. Limited by short target sequences, we did not compare relative efficiencies for the different systems, which would require a large scale study involving multiple targets for each Cas protein. For example, it remains to be determined if Type V-A Cpf1 provides an advantage in promoting HDR in streptomycetes (Jiang, *et al*, 2017) compared to Type II Cas proteins since fnCpf1 and saCas9 can achieve the same editing efficiency in the cases that we have tested.

The collection of CRISPR-Cas systems that have been characterized and harnessed for various applications has rapidly expanded (Cong, *et al*, 2013; Zetsche, *et al*, 2017), but it remains a challenge to edit the genomes of most bacteria due to the lack of robust HDR and NHEJ. In *E. coli*, this is circumvented by introducing additional phage recombineering machinery (Ronda, *et al*, 2016; Reisch, *et al*, 2015), which may not be applicable in other bacteria. In organisms where the spCas9 system has failed due to low efficiency or high toxicity, the use of alternative CRISPR-Cas systems has been successful (Adli, 2018; Rock, *et al*, 2017; Sun, *et al*, 2018). Therefore, until we can better understand the mechanisms and rules for successful precise editing by different CRISPR-Cas systems (Chakrabarti, *et al*, 2018), having a diverse selection of functional Cas proteins provides us with the versatility to overcome the limitations of individual systems. Beyond genome editing, multiplexing with orthogonal catalytically inactive Cas proteins with different guide specificities may allow transcriptional control of different gene subsets in actinomycetes using CRISPR interference (Rock, *et al*, 2017; Tong, *et al*, 2015).

## Acknowledgments

This work is supported by the Agency for Science, Technology and Research (A*STAR), Singapore and National Research Foundation, Singapore [NRF2013-THE001-094 to MMZ, FTW, YHL] and the A*STAR Visiting Investigator Program [HZ]. The authors declare no conflict of interest.

## Supplementary data

**Supplementary Table 1.**
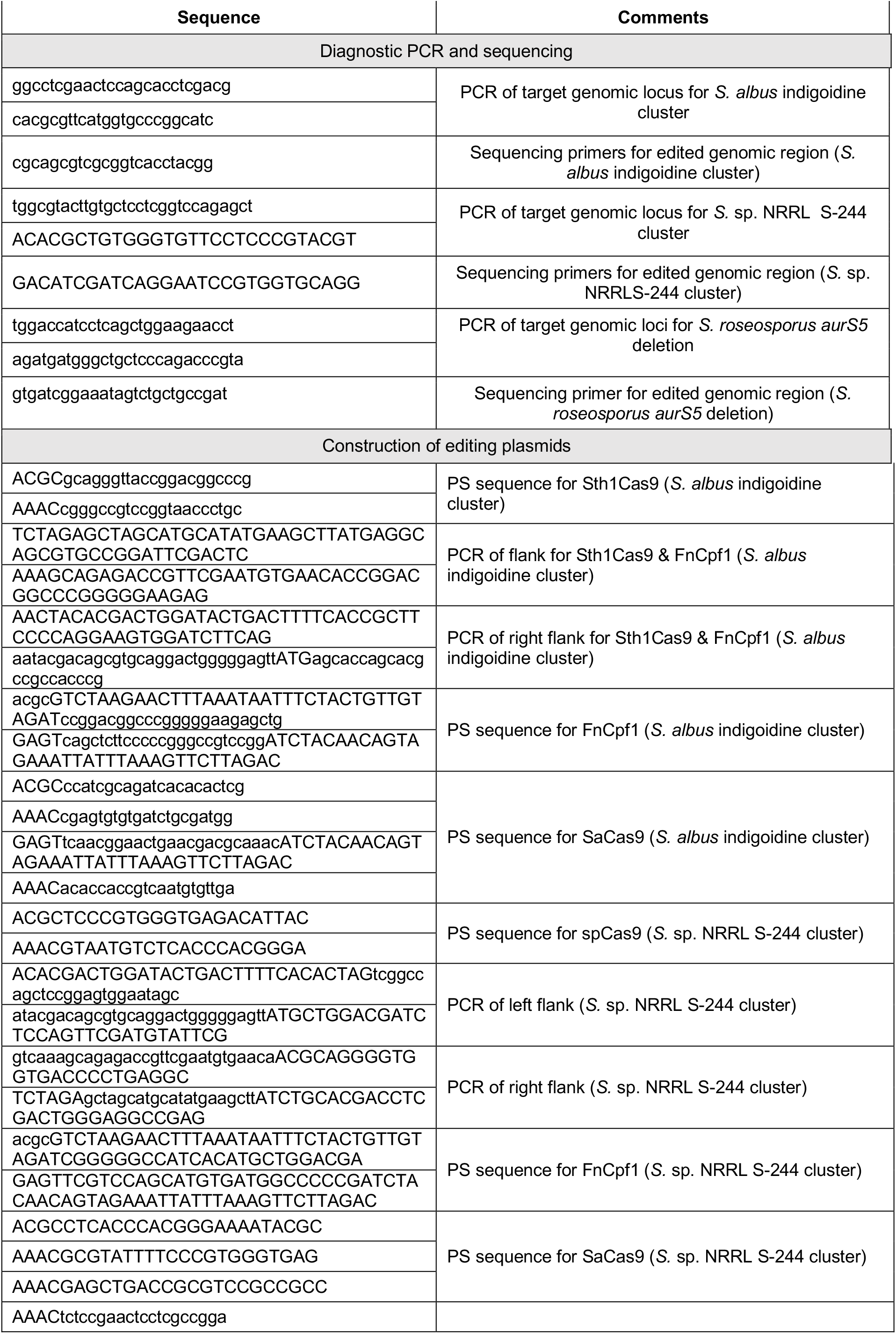
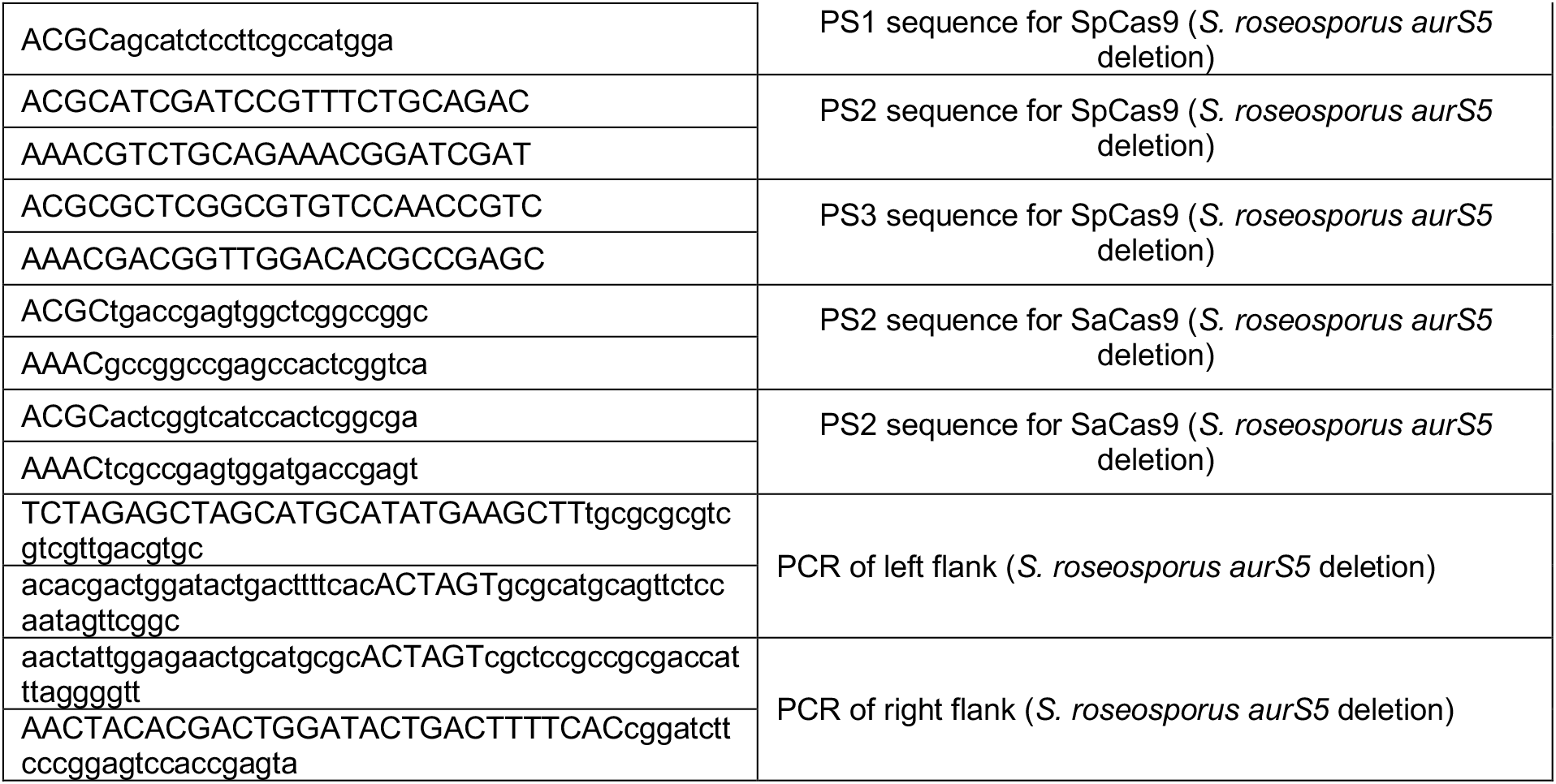
Oligonucleotides used in this study.

**Figure S1.**
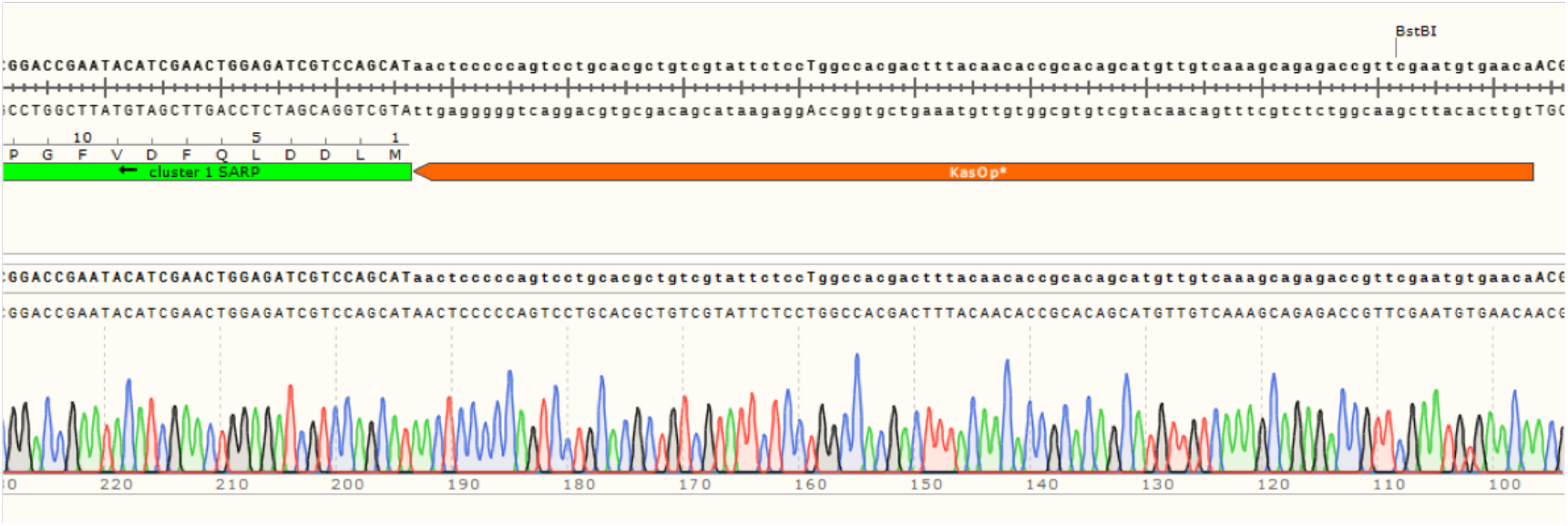
Representative trace of edited genome sequence (insertion of *kasO*\*p) for *Streptomyces* sp. NRRL-244.

**Figure S2.**
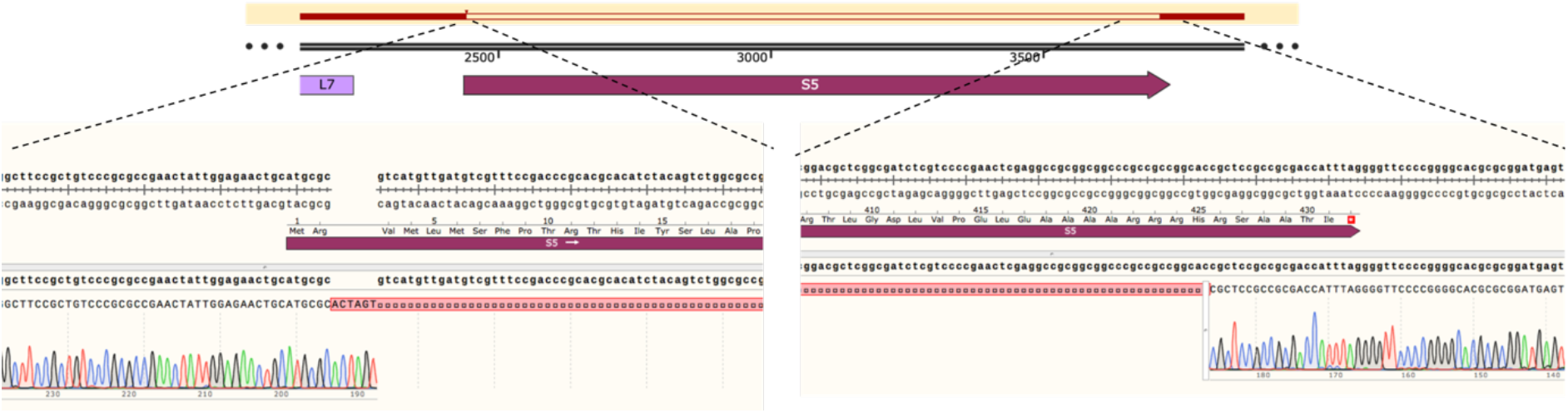
Representative trace of edited genome sequence (deletion of *aurS5*) for *Streptomyces roseosporus*

